# Large scale active-learning-guided exploration to maximize cell-free production

**DOI:** 10.1101/751669

**Authors:** Olivier Borkowski, Mathilde Koch, Agnès Zettor, Amir Pandi, Angelo Cardoso Batista, Paul Soudier, Jean-Loup Faulon

## Abstract

Lysate-based cell-free systems have become a major platform to study gene expression but batch-to-batch variation makes protein production difficult to predict. Here we describe an active learning approach to explore a combinatorial space of ~4,000,000 cell-free compositions, maximizing protein production and identifying critical parameters involved in cell-free productivity. We also provide a one-step-method to achieve high quality predictions for protein production using minimal experimental effort regardless of the lysate quality.

## Main text

Cell-free systems, especially lysate-based systems, are major platforms for both prototyping of genetic circuits and understanding of fundamental processes ^1–7^. They provide fast gene expression kinetics, low reaction volumes, allowing high-throughput measurements and simplified gene characterization via decoupling protein production from host physiology ^8–12^. Cell-free systems could disseminate among laboratories and be standard methods for molecular biology if efficient and predictable protein productions were guaranteed. Ribosomes, native polymerases and cofactors concentrations remain arduous to control as they are provided by the lysate ^13,14^, making the efficiency of cell-free systems variable. A great challenge is to develop a lysate-specific optimization method for cell-free composition to maximize protein production. Using a Design of Experiment approach, Caschera *et al*.^13^ explored cell-free compositions by varying one compound concentration at a time and obtained a 10 fold increase of protein production in a lysate-based cell-free system. Such results reveal the considerable margins of improvement of protein expression in such systems.

Here, we use an active learning approach^15,16^ to explore, optimize and understand the impact of cell-free composition on protein production in cell-free systems. We demonstrate that sufficient amount of data can be obtained to train a machine learning algorithm, achieve high quality predictions and increase protein production by 34 times. We next show that only 20 informative compositions are enough to train our machine learning model and obtain accurate predictions. This approach enables to maximize protein production on different cell lysates with minimal experimental effort.

To study cell-free systems productivity, we developed an automatable strategy coupling an acoustic liquid handling robot (Echo 550, Labcyte, USA) and a plate reader (Infinite MF500, Tecan, USA) to measure ~4000 cell-free reactions (including controls and triplicates) and provide data to train a machine learning model. The lysate was obtained by sonication and supplemented with compounds described in Fig. 1a. The reference concentrations is based on the protocol developed by the Noireaux laboratory ^17^ (see methods, Supplementary Fig. 1). We fixed 4 concentration levels for each of the 11 compounds leading to a combinatorial space of 4,194,304 possible compositions (Fig. 1a). Protein production was measured using the fluorescence level from the expression of *sfgfp* under control of a constitutive promoter (Fig. 1b, Supplementary Table. 1). In order to compare measurements between plates, we maximized a relative fluorescence level named yield hereafter (Fig. 1b). The yield is defined as the ratio of the fluorescence produced with a chosen composition divided by the fluorescence obtained with the reference composition (Fig. 1b). To explore our vast combinatorial space, we used an active learning strategy ^15^, combining both exploration and exploitation to increase the yield and reduce model uncertainty (Fig. 1c). Each iteration started with 102 new cell-free compositions to be tested. The fluorescence level was measured in a plate reader and fed to an ensemble of neural networks (Fig. 1c, see methods). Our active learning loop was initiated with a training set of 102 cell-free compositions (see methods: 22 chosen and 80 random compositions, Fig. 1c). The first iteration already led to a maximum of 10 fold improvement of the yield (Fig. 1d). As expected, the prediction accuracy was very low (Fig. 1e). After 7 iterations, we reached a maximum for both the yield (Fig. 1d) and the prediction accuracy (Fig. 1e). Eventually, we stopped at 10 iterations as we were not able to increase neither the yield nor the prediction accuracy of our model (Fig. 1d, Fig. 1e, see methods). Throughout our workflow, we measured fluorescence levels in 1017 cell-free compositions and validated the efficiency of our method with a high quality predictions score (R^2^=0.93) and a maximum of 34 fold increase of the yield. The 1017 cell-free compositions were sorted, from low to high yields, to observe the relationship between yield and composition (Fig. 1f). An increase of Mg-glutamate, K-glutamate, Amino Acids and NTPs concentrations and a decrease of cAMP, spermidine and 3-PGA concentrations can be noticed with increasing yield (Fig. 1f). We used a mutual information analysis (see methods) to reveal the dependence between our 11 compounds concentration and the yield. Mg-glutamate, K-glutamate, Amino Acids, cAMP, spermidine, 3-PGA and NTPs exhibit a score between 0.25 and 0.75, confirming that a variation of their concentrations strongly impacts protein production (Fig. 1g). Variation of tRNA, CoA, NAD and Folinic Acid concentrations have little impact on the yield (Fig. 1g).

**Fig. 1.**
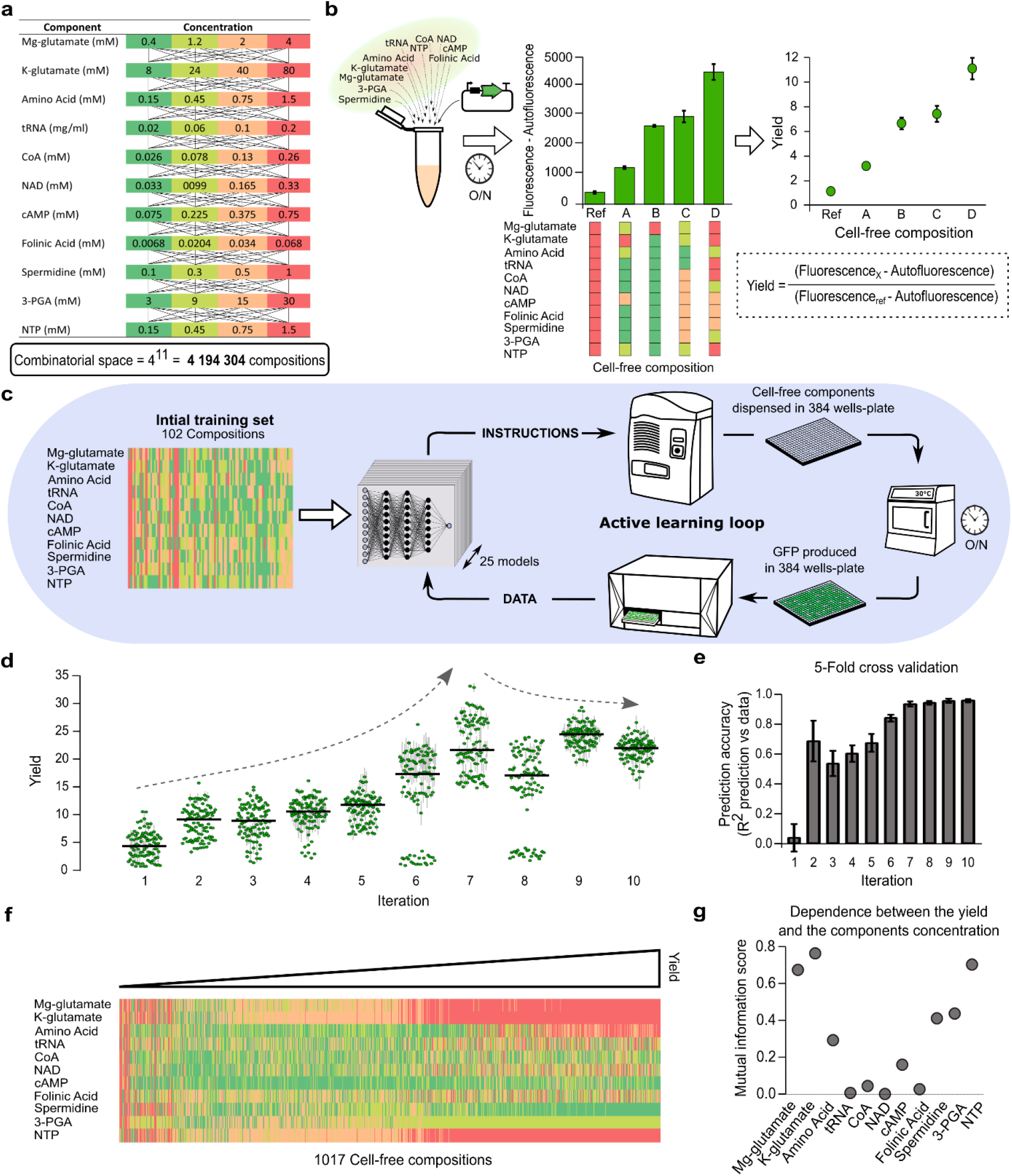
Active learning loop to explore the composition of a cell-free system. **a**, List of chemicals added to the cell-free mix in addition to PEG-8000, HEPES and the lysate. Four concentrations have been chosen for each chemical. The concentration in red is the highest concentration, then orange, light green, dark green stand for 50%, 30% and 10% of the highest concentration. **b**, An example of fluorescence obtained using 4 cell-free compositions with our plasmid (10 nM). The autofluorescence value is measured with the reference composition without DNA and subtracted from every measurement in the plate. The yield is the ratio between the fluorescences of a composition x and the reference composition. **c**, Illustration of the active learning approach used to explore the combinatorial space of cell-free composition and trained an ensemble of 25 machine learning models. **d**, Yield evolution amongst 10 iterations. The green dots are the mean yields of 3 replicates obtained in the same plate with the same composition. The vertical grey lines stand for the standard deviation of the 3 replicates. The horizontal black line is the median value of all the yields obtained during an iteration. The Arrows represent the evolution of the maximal yields value. **e**, Quantification of the predictive accuracy of the model using a 5-Fold cross validation. **f**, Cell-free compositions tested in the study sorted by yield level. A row stand for one mix composition, the colour code is the same as in panel a. **g**, Results of a mutual information analysis, using the 1017 compositions, of the relationship between the yield and each chemical compound.

Next, we investigated whether protein production in cell-free using lysates made in other conditions (different experimentalists, using a different strain or supplemented with antibiotics) could be quickly predicted with a one-step method (Fig. 2a). We selected 102 cell-free compositions representative of the 1017 already tested with the original lysate (see methods, Supplementary Fig. 2a). Amongst the 102 compositions, 20 were used to train the model and 82 to test its predictive accuracy (Fig. 2a). The challenge lies in the model’s ability to accurately predict a large diversity of yields based on a small training dataset. The 20 compositions (magenta dots in Fig. 2) were chosen to be highly informative (see methods, Supplementary Fig. 2b). We used the same 20 and 82 compositions to train and test our model with all the lysates used in Fig 2. With new lysates prepared by other experimentalists (labeled lysate_PS and lysate_AB), similar cell-free compositions led to different yields but the compounds exhibiting a high impact on protein production remained the same (Fig 1g and Supplementary Fig. 3). The maximum yield amongst the 102 tested compositions differs from one lysate to another, with a maximum yield at 23 and 26 for the lysate_PS and lysate_AB respectively (Fig. 2a,b). The 102 yields obtained with the original lysate, labeled Lysate_ORI, are presented in Supplementary Fig. 4a. The yield used previously is a relative measurement (Fig. 2 and Supplementary Fig. 5) which does not allow absolute comparison between our cell-free systems. We calculated a global yield (calculated with the Lysate_ORI as a global reference, Supplementary Fig. 4b) and observed a maximum global yield 1.5 times higher with lysate_PS than lysate_AB (Supplementary Fig. 4c). These results highlight the variability in lysates quality even when they are prepared in the same laboratory with the same strain and protocol. Despite these differences, we achieved high quality predictions with both lysates (Fig. 2 a,b). We obtained a R^2^ ~ 0.9 for both lysates and linear fits with intercepts of 0.2 / 0.1 and slopes of 0.8 / 1.01 with lysate_PS, lysate_AB respectively (Fig. 2 a,b). These results validate our approach to both maximize protein production and accurately predict protein production regardless of the experimentalists who prepared the lysate.

**Fig. 2.**
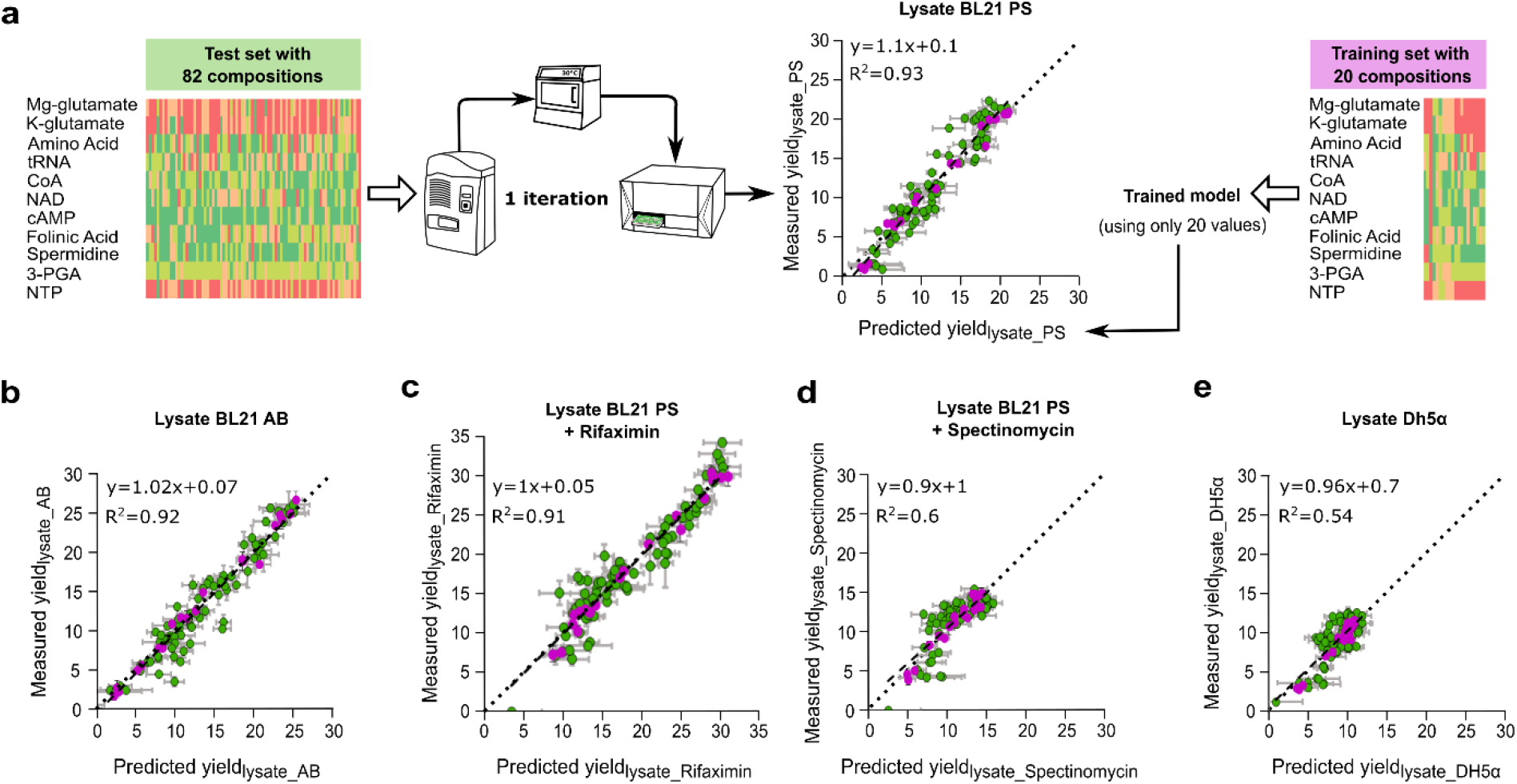
One-step method to predict protein yield in cell-free systems. **a**, Illustration of the method used to predict the yield of protein expression with a new lysate, labelled PS, made by another experimentalist. The training of the model is based on yield measurements of 20 compositions (Magenta circles). The choice of the 20 combinations leading to the best predictions is described in the methods. The yield obtained with 82 compositions (Green circles) were measured and compared to the model predictions to test its accuracy (R^2^ value). The yield is specific to each lysate as the reference composition used the same chemicals concentration as in Fig. 1 but with different lysate. **b**, Comparison of the yields obtained with the lysate AB (made by a third experimentalist) vs the model predictions. **c**, Comparison of the yields obtained with the lysate, of panel a, supplemented with 0.25 mg/mL of rifaximin vs the model predictions. **d**, Comparison of the yields obtained with the lysate, of panel a, supplemented with 0.5 mg/mL of spectinomycin vs the model predictions. **e**, Comparison of the yields obtained with a lysate obtained from the stain DH5a vs the model predictions. In all panels, the model predictions are based on a model trained with the same 20 compositions and the same test set of 82 compositions only lysate differs. In all panels, the horizontal grey lines stand for the standard deviation of 3 replicates. The vertical grey lines stand for the standard deviation of 25 predictions.

We then challenged our method by interfering with the transcription or translation processes to mimic lysates of lower quality. By adding rifaximin (Fig. 2c) or spectinomycin (Fig. 2d) to the cell-free mix, we interfered with the transcription or translation apparatuses respectively. The two antibiotics led to a strong decrease in absolute protein production (Supplementary Fig. 4c) but opposite behaviours can be observed (high versus low room for yield improvement, Fig. 2 c,d, Supplementary Fig. 5c,d). When the transcription process is impaired, we obtained a prediction of high accuracy with a R^2^ of 0.91 and linear fit intercepts of 0.2 and slopes of 0.9 (Fig. 2c). The cell-free containing rifaximin exhibits a high leeway for yield improvement (Fig. 2c and Supplementary Fig. 5c) with a maximum yield of 35 amongst the 102 cell-free compositions. When the translation process is impaired, the yield is capped to a maximum improvement of 15 (Fig. 2d, Supplementary Fig. 5d). The R^2^ value observed in Fig. 2d is lower but the linear fit exhibit an intercept of 0.1 and slopes of

0.9. Thus, we obtained accurate prediction for the low and high yields value but the intermediate yields remains difficult to estimate. Such predictions stay powerful to maximize protein production as extreme values are captured and provide precious information concerning the lysate quality (Supplementary Fig. 6, **supplementary note 1**).

Eventually, we tested our method with a lysate prepared using the strain DH5α. As observed with the lysate supplemented with spectinomycin, the R^2^ value is low but the linear fit of the data exhibits an intercept of 0.07 and slopes of 0.96. The maximum global yield obtained with this lysate was low, as expected for a strain not optimized for protein production^18^ (Supplementary Fig. 4c). Nevertheless, with half of the tested cell-free compositions, the Lysate_DH5α-based cell-free exhibits a high global yield (Supplementary Fig. 4c). The yield exhibits a similar behaviour as the lysate supplemented with spectinomycin, suggesting an impaired translation process (little room for yield improvement, Fig. 2d,e, Supplementary Fig. 5d,e), but with a higher level of protein production.

Our method enables a fast lysate-specific optimization of the cell-free composition to predict and maximize protein production (Fig. 2, Supplementary Fig. 7, and **Supplementary note 2**). Our results suggest that the optimization of the cell-free composition mainly improves the efficiency of the translation apparatus as we observed a limited improvement with an impaired translation. On the contrary, a damaged transcription machinery can be balanced by the optimization of the cell-free composition. Our approach gives precious information about the room for protein production improvement of a home-made cell-free system, the impact of each compound on cell-free productivity and the efficiency of the transcription and translation processes. Our method, based on the measurements of GFP production with the same 20 cell-free compositions used in this work to train the model provided, can be easily extended to any other bacterial-based cell-free ^4,19-20^ to investigate cell-free optimisation beyond *E. coli* cell-free systems. As our model is not based on mechanistic hypotheses, our method can be extended to cell-free systems using other organisms as yeast, insect, plant or human cells after performing new explorations to find the 20 most informative compositions for those cell-systems.

## Acknowledgements

O.B is supported by Genopole ‘Allocation Recherche 2017’ and CRI Paris ‘Short-term Fellows’. M.K is supported by DGA (French Ministry of Defense) and Ecole Polytechnique. PS is supported by ANR (grant number ANR-18-CE33-0015). A.C.B acknowledge funding provided by the ANR SINAPUV grant number ANR-17-CE07-0046. A.P is supported by INRA (National Institute for Agricultural Research) and an idEx interdisciplinary scholarship from the University of Paris-Saclay. J-L. F acknowledges support from BBSRC/EPSRC (grant number BB/M017702/1). The chemogenomic and Biological Screening Platform of Pasteur is funded by the Global Care initiative and Institut Carnot Pasteur MS. We thank the Faulon Lab and Agou Lab members for fruitful discussions. We thank Claire Donnat (from Stanford University) for the data analysis advice that she kindly gave us. We thank Stephen McGovern (from INRA) who generously provided purified sfGFP.

## Author contributions

O.B, M.K and J-L F designed experiments. O.B performed experiments. M.K developed and performed model simulations and liquid handler programming. O.B and M.K performed data analysis. O.B and A.Z collected data. O.B, A.P, A.C.B and P.S provided lysates. A.P cloned and maxi prep the plasmid. O.B, M.K, A.Z and J-L F wrote the paper. All authors approved the manuscript.

### Competing interests

The authors declare no competing interests.

## Code availability

Code is available on GitHub at https://github.com/brsynth.

## Methods

### Bacterial strains and DNA constructs

Strains BL21 DE3 (B F– ompT gal dcm lon hsdSB(rB–mB–) λ(DE3 [lacI lacUV5-T7p07 ind1 sam7 nin5]) [malB+]K-12(λS)) and DH5α (F– endA1 glnV44 thi-1 recA1 relA1 gyrA96 deoR nupG purB20 φ80dlacZΔM15 Δ(lacZYA-argF)U169, hsdR17(rK–mK+), λ–) were used to prepare the different lysates in this study. Our *sfgfp* plasmid was obtained by modification of the RBS of the plasmid pBEAST-J23101-sfGFP^9^. We used PCR amplifications using the reverse primer GCGGTCTCACATCTACTATTTCTCCTCTTTCTCTACTAGCTAGC and foward primer GCGGTCTCAGCTTACTTTATCTGAGAATAGTC with the backbone, and reverse primer p CCGGTCTCAAAGCTTATCATCATTTGTACAGTTCATCC and GCGGTCTCAGATGCGTAAAGGCGAAGAG foward primer with the *sfgp* sequence. The PCR amplifications was followed by a golden gate assembly using BsaI and T4 ligase (New England Biolab) and transformed into chemically competent *E. coli* top10.

### Plasmid preparation

We noticed with preliminary experiments that the same cell-compositions gave different results when we used plasmid DNA from miniprep done on different days using the same kit. The whole project was done using aliquots from the same initial batch of *sfgfp* plasmid. The plasmid was extracted from a 600 ml LB of *E. coli* top 10 using the Plasmid DNA purification NucleoBond Xtra Maxi of Macherey-Nagel. The 500 µl aliquots were stored at −80°C. The whole project was done using aliquots from the same initial batch of DNA. The final *sfgfp* plasmid concentration in every reaction was 10nM.

### Cell-free reagents preparation

As the reagents preparation can have a significant impact on cell-free efficiency^21^, all our reagents except spermidine and Mg-glutamate (we run out of those two compounds during the study) came from aliquots of the same initial batch. We did not see an impact on our control when the spermidine and Mg-glutamate were renewed.

### Cell lysate mix preparation and reactions

The cell lysate preparation is based on the protocol of Sun *et al.*^17^. Briefly, the protocol of Sun *et al.*^17^ is a 5 day protocol in three phases: harvest cells (colonies grow on plate overnight at 37°C, 50 ml preculture at 37°C for 8 h, 12 liters of cultures at 37°C until OD_600_= 1.5), lysate preparation (multiple pellet washing followed by beads-beating to obtain an lysate). The protocol was modified by using sonication instead of use of a bead beater to obtain BL21 or DH5α cell lysates. After washing the cells as following the Sun *et al*. protocol (Day 3 step 18) with S30A buffer (14 mM Mg-glutamate, 60 mM K-glutamate, 50 mM Tris, 2 mM DTT, pH 7.7), the cells were centrifuged 2000×g for 8 min at 4 °C. The pellet was re-suspended in S30A (pellet mass (g) × 0.9 ml). The solution was split in 1.5 ml aliquots in 2 ml Eppendorf tubes. Eppendorf tubes were placed in a cold block and sonicated using Vibracell 72408 (from Bioblock scientific) using the following procedure:20 s ON—1 min OFF—20 s On—1 min OFF—20 s ON. Output frequency 20 kHz, amplitude 20%.The remaining protocol followed the procedure of the Sun *et al.* protocol for day 3, step 37. mRNA and protein synthesis are performed by the molecular machinery present in the lysate, with no addition of external enzymes. Reactions take place in 10.5 μL volumes at 30 °C in 384-well plate. Note that we kept the 50 mM HEPES and 2% PEG-8000 fixed in every reaction. Lysate_ORI, Lysate_PS and Lysate_AB were obtained from the same *E. coli* strain BL21 in the same laboratory with the same sonicator and centrifuge. The Lysate_ORI came from one-batch prepared from 12 Liters of BL21 culture. The 12 liters culture were separated in 4 Liters culture. The culture were inoculated, grown and their pellets were washed on different 3 days then freeze and stock at −80°C. Then, the pellets were weighed, resuspended in S30 buffer, pooled, sonicated, centrifuged, mixed and aliquoted on an extra day. The Lysate Lysate_PS, Lysate_AB and Lysate_PS and Lysate_DH5α were each obtained from 2 liters culture. For the Lysate_spectinomycin and Lysat_rifaximin, the final concentration of rifaximin and spectinomycin were 0.25 mg/ml and 0.5 mg/mL respectively. They were added to the cell-free reactions using Lysate_PS.

### sfGFP purification

The sfGFP was produced in *E. coli* culture. After a 10 min centrifuge at 4000g, the pellet was resuspended in 20 mM Tris (Ph8), 0.2 M NaCl and sonicated (Output frequency 20 kHz, amplitude 40% with the Vibracell 72408). After sonication, the solution was centrifuged (4000g, 15 min). The proteins in the supernatant were purified and fractionated using ammonium sulfate. The sfGFP was isolated at more than 70% saturation. The solution was centrifuged (4000g, 15 min) and the pellet resuspended in 20 mM Tris (Ph8), 100 mM NaCl. The solution was dialysed overnight in 20 mM Tris (Ph8), 100 mM NaCl. Eventually, for the last step of purification, we used a Mono Q anion exchange chromatography column (GE Healthcare) and obtained a solution of 90% sfGFP. The final solution dialyse in a solution 0 mM Tris (Ph8), 100 mM NaCl and 50% glycerol leading to a final concentration of 7.62 mg/ml. To obtain an absolute quantification of the protein production in cell-free, we measured the sgGFP fluorescence in wells containing 10.5 µl of sfGFP solution at different concentration. The G-yield values are calculated as described in Supplementary Fig. 4b with the fluorescence measured from sfGFP and no autofluorescence divided by the cell-free mix lysate_ORI autofluorescence and the reference fluorescence obtained from our plasmid in a cell-free mix with lysate_ORI.

### myTXTL commercial kit

We used the commercial kit: myTXTL from Arbor Biosciences (Sigma 70 Master Mix Kit, (USA). We used both our plasmid (10 nM final concentration) and the control plasmid, pTXTL-P70a(2)-deGFP (20 nM final concentration) provided by Arbor Biosciences. The 2 plasmids were expressed with the reactions provided with myTXTL kit and with the optimised cell-free reaction with the Lysate_ORI Supplementary Fig. 7b.

### Fluorescence quantification

We used a plate reader Infinite MF500 (Tecan) to measure fluorescence in 384-well plates (Nunc 384-well optical bottom plates, Thermo-Scientific). The excitation wavelength was fixed at 425 nM, the emission at 510 nM and the gain at 50. We measured 5 fluorescences values for each well as a quality control of the plate reader measurements. The fluorescence was measured from the top of the 384-well plates with no lid.

### Echo liquid handler

We used the Echo software Cherry Pick to program the Echo 550 liquid handler. The software was programmed using CSV (comma separated values) files that gave machine-readable instructions: namely the well it had to take liquid from (containing pure reagents), the well the liquid was destined to and the volume that was to be taken. It allows us to program the content of each individual well separately. We calculated the concentrations of our chemical compounds stocks so the final volumes sent to the destination well were multiples of 2.5 nL (the droplet size managed by the Echo machine). The scripts generating the CSV file are presented below in *concentrations to instructions workflow*. We chose our stock volumes so that the minimal volume to transfer was 12.5 (=5 droplets). The chemical compounds were dispensed using BP2 fluid class except for K-glutamate and 3PGA (CP fluid class).

### General script descriptions

All scripts mentioned below were written in Python (version 3.6.5), executed in Jupyter notebooks (version 1.0.0). Scripts are available online at github (https://github.com/brsynth). The libraries numpy and csv were used to handle files between different scripts. We used scitkit-learn^22^ (version 0.19.1) for all model training.

### Concentrations to instructions workflow

The details of those scripts are described in the READme file of the ECHO_handling_scripts of our code. Roughly, it proceeds in 4 steps:

- *Complete concentrations*: Taking as input a file containing only concentrations of interest for the machine learning algorithm, it adds information of values that are constant across all conditions, such as the lysate quantity.
- *Concentration to volume:* This file converts a csv file concentrations-to a file of volumes one wants to test (in triplicates). This is due to the fact that the ECHO liquid handler needs volumes as inputs.
- *Optional*: we sorted those volume files according to water content. This allows us later to manually pipet important water volumes so that the robot only adjusts small volumes.
- *Volume to echo:* This file converts a set of transfer volume quantities to the csv file expected by the ECHO liquid handler (instructions files). It also provides a file containing the name of the wells with their corresponding transfer volumes. This file is used to match the well compositions with the fluorescence measurements obtained later with the plate reader. The amino acids and water were pipetted manually (for volumes > 1µl).
- *Named volumes to concentrations:* maps the volumes and the associated well name to a concentration file with the associated well name, for integration with the fluorescent plate reader at the next step.

The script matched the named concentration with the yield value as described in *Data analysis* of those methods.

### Data analysis

We provide a script to map the fluorescence quantification (see *fluorescence quantification* above) to the tested concentrations with well names (last step of *concentrations to instructions workflow* above). We performed outlier removal based on the following criteria: if the coefficient of variation, amongst 3 replicates, was higher than 30%, we removed the value farthest from the other 2. This concerned 27 values of our 1017 values tested during the active learning. Those are identified in the online data on Github with the third value of fluorescence is set to −1. This script also outputs csv files allowing for visualisation of where the outliers are, in order to spot potential border effects. It also separately outputs the outliers for further analysis.

### Data normalisation

We normalised using the following equation:

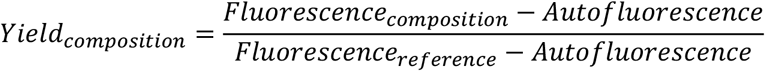

Where autofluorescence is the fluorescence measured in the cell-free reaction supplemented with water and using the reference composition. The yield exhibited in Fig. 2 used a cell-free reaction with the new lysate to measure the autofluorescence and the fluorescence with the reference composition. In supplementary Fig. 4, all the yields are calculated with autofluorescence and reference fluorescence of the Lysate_ORI.

### Quality controls

In every 384-well plates we measured 13 control compositions (in triplicate) including the reference composition with and without DNA In each 384-well plate, we used 2 rows of controls: A and P. The controls in row A never changes. The controls in row P changed throughout the workflow. We used the compositions leading to higher yields in the previous iteration. When analysing our controls, we checked whether the yields were identical from plate to plate (R² > 0.75 between new plate and all previous plates on yield of controls). Plates with R² >0.75 when compared to all previous plates, or systematically above or below other plates are discarded and the same combinations were tested again.

### Initiation of the Machine learning

For the first plate of the active learning, we proceeded as follows. We chose 22 concentrations that we wished to test: fixing all reagents at the maximum allowed concentration, except one which was at the lowest (11 combinations) and fixing all reagents at the minimum allowed concentration except one which was at the highest (11 combinations). The rest (80 compositions) was filled randomly.

### Model training

The models were trained as follows.

- Input data is normalised: each component maximum concentration is 1, and the other values take discrete values of 0.1, 0.3, or 0.5 as described in the legend of Fig. 1. While unnormalised inputs could be used, we strongly encourage normalisation due to scale differences between the inputs.
- We train an ensemble of n models, where n = 25. For each model, we train it 10 times (models_number) using the whole dataset at the moment (e.g. 3×102 values at the 3rd iteration). Training the model multiple times allows for optimising for random weight initialization of the model. We keep the best model (with the highest regression from scitkit-learn R^2^ score).
- Multilayer perceptrons give the best results (random forests and linear regressions were also investigated early on). They are trained with the default parameters from scitkit-learn except the following parameters: maximum iteration of 20000, adaptive learning rate, adam solver, early stopping and the following layers: (10, 100,100, 20)
- We obtain mean and standard error from our predictions by taking the mean and standard error from the n results generated by our ensemble of n models.

### Active learning

The workflow used the data from all the available plates as an input. It trains an ensemble of 25 models and returns instructions for the following round. Here is the detailed process:

- For N times (N= 100,000):

- Randomly sample a composition in the composition space (Fig. 1a)
- If a composition was drawn previously (either in a previous experiment or during current selection), neglect it.
- Predict mean and standard deviation for all 100,000 points using the ensemble of 25 models previously trained.
- Select the best set of compositions, according to the following Upper Confidence Bound (UCB) formula: exploitation * yield_pred_mean + exploration * yield_pred_std, with exploitation = 1 and exploration = 1.41. Our scripts output the best 500 compositions based on the mean and std predictions of the yields. A high std value stands for an uncertain yield value. We output compositions for full exploitation, full exploration and maximisation of the above formula but use the third option for the rest of the workflow. We are therefore querying points with both high yield and uncertainty.

### Model statistics

For model statistics presented in Fig. 1e, we used the same models as described in the active learning section above, but using 5 fold cross validation instead of the whole training set. The full dataset is separated into 5 subsets then the 25 models are trained on 4 subsets, and used to predict the 5th, where scores are obtained. This is done 5 times, once on each subset. The scores presented in Fig. 1e are the mean and standard deviation of those 5 scores.

### Mutual information calculation

Mutual information is a method to quantify the mutual dependence between two variables. This concept is intrinsically linked to the concept of entropy and is especially useful to quantify non-linear relationships between variables. More information on the theory behind this method can be found in the review ‘estimation of mutual information^23^ and in sci-kit learn documentation^22^. It was calculated using the feature_selection.mutual_info_regression function from scitkit-learn^22^ (version 0.19.1) between each feature and the yield (compounds_effect_analysis/mutual_information_analysis jupyter notebook) with default parameters.

### Identification of informative points

To identify the most informative points, we proceeded in the following manner:

We did 1000 iterations of the following procedure:

- Randomly sample n combinations from the dataset (n=20 out of a dataset of 102 values for Fig. 2

- Train models on those points using the strategy presented in model training for each lysate
- Predict on the other points (82 for Fig. 2) for each lysate
- Obtain the average score on all lysate,-s
- Keep those combinations if this average is better Note: Data is saved every 100 iterations

### Maximization of the protein production for future users

Users must do the following experiments:

- Maxiprep a LB culture of our plasmid (or MyTXTL plasmid)
- Measure the yields (or absolute fluorescences) in the 20 cell-free compositions described in Fig. 2a

Then, in order to apply our method to a new extract, a jupyter notebook called predict_for_new_lysate is available. It takes as input a csv file containing the 20 tested concentrations and the 20 corresponding yields and standard error values. It provides as an output a file to maximise exploration, exploitation or a combination of both as in the active learning loop. For obtaining the highest possible yield, it is recommended to take the exploitation results, which contain the highest predicted yields. It must be noticed that several cell-free compositions can be predicted to reach maximum yield or values in the same range. The algorithm provides mean yields value with standard deviation errors and so several yields will be equivalent to the maximum value. During this study we provided yield values to our training algorithms but absolute fluorescence can also be used if a user does not need to compare fluorescence values measured on different 384-well plates.

## Supplementary information

### SUPPLEMENTARY NOTE 1 Deterministic model of protein production behavior in cell-free system with an imparied translation process (Supplementary Figure 6)

#### Assumption 1

Adding spectinomycin lead to a similar impact on the translation process as a decrease in concentration of the available ribosome. Spectinomycin binds to the 30S subunit stopping protein synthesis. Thus, a subset of ribosomes should be unavailable for translation.

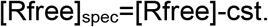

#### Assumption 2

We simplified our calculation by considering that a variation in cell-free composition has a similar impact on both V_max_ and K_M_.

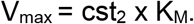

#### Assumption 3

The relationship of transcription efficiencies (noted TxE) between lysates is modeled by a linear relationship with a negligible intercept. We observed such a linear relationship (with an intercept close to 0) between yields from lysates with and without a damage transcription machinery in Supplementary Figure 5c.

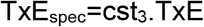

#### Assumption 4

The variation in cell-free composition mainly impact the translation process. We observed in Supplementary Figure 5d that a lysate with a damaged translation machinery is poorly improved by a change in cell-free composition. The opposite is observed with a damaged transcription machinery in Supplementary Figure 5c suggesting that the efficiency of the translation machinery is the limiting factor for cell-free improvement and not the efficiency of the transcription machinery.

TxE=cst_4_ (TxE is independent of the variations in cell-free compositions)

We used the well-defined model of the translation efficiency (TlE) based on a Michaelis-Menten equation^24^:

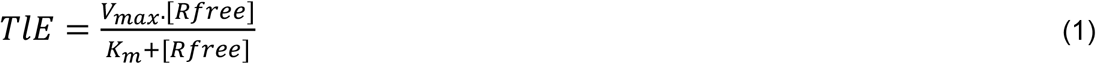

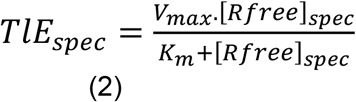

where V_max_ and K_M_ values depends on the RBS sequence and the cell-free composition. [Rfree] stands for the concentration in available ribosomes.

Assumption 1: [Rfree]_spec_=[Rfree]-cst.

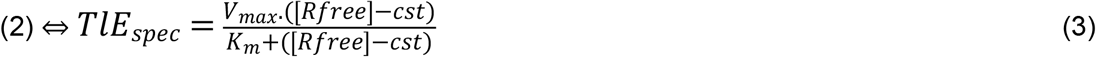

Assumption 2: V_max_ = cst_2_ × K_M_

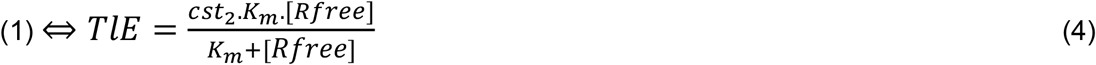

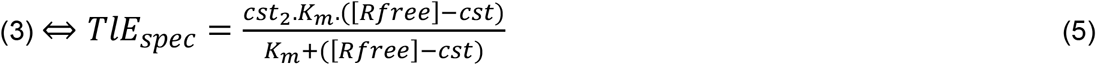

Thus,

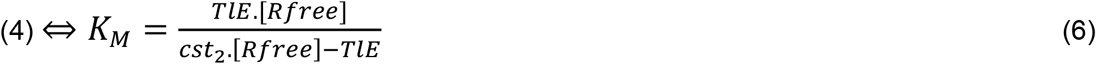

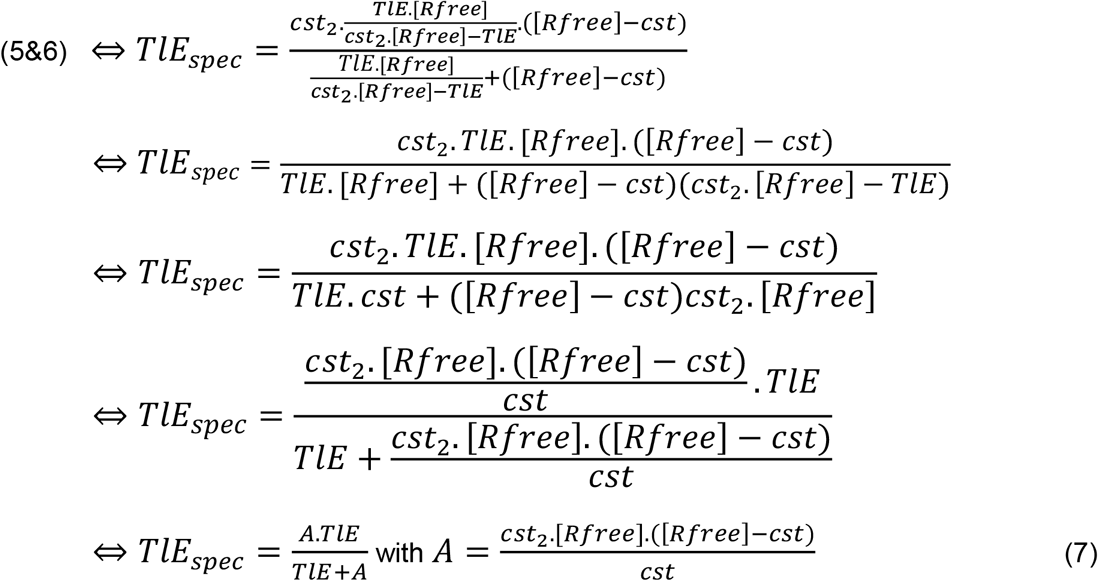

The protein production (and so the yield) is the result of the expression of *sfgfp* by the transcription and translation processes.

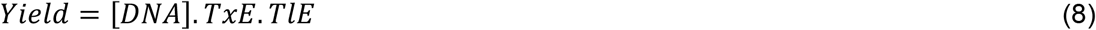

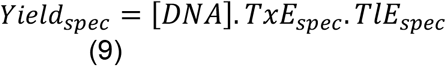

Assumption 3: TxE_spec_=cst_3_.TxE. Moreover, the DNA concentration is the same in every cell-free reaction so [DNA] =cst_5_.

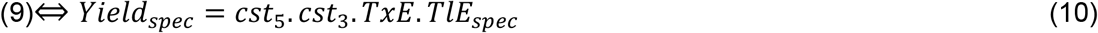

Assumption 4: TxE=cst_4_

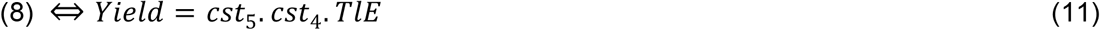

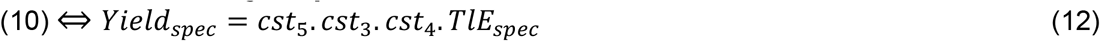

Then,

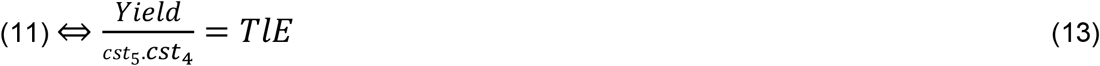

and

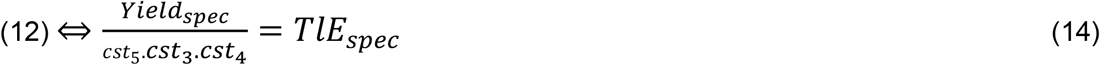

Then,

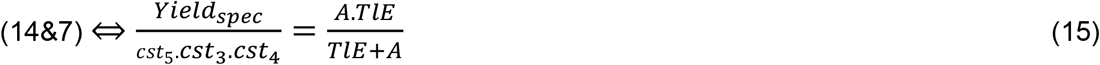

Then,

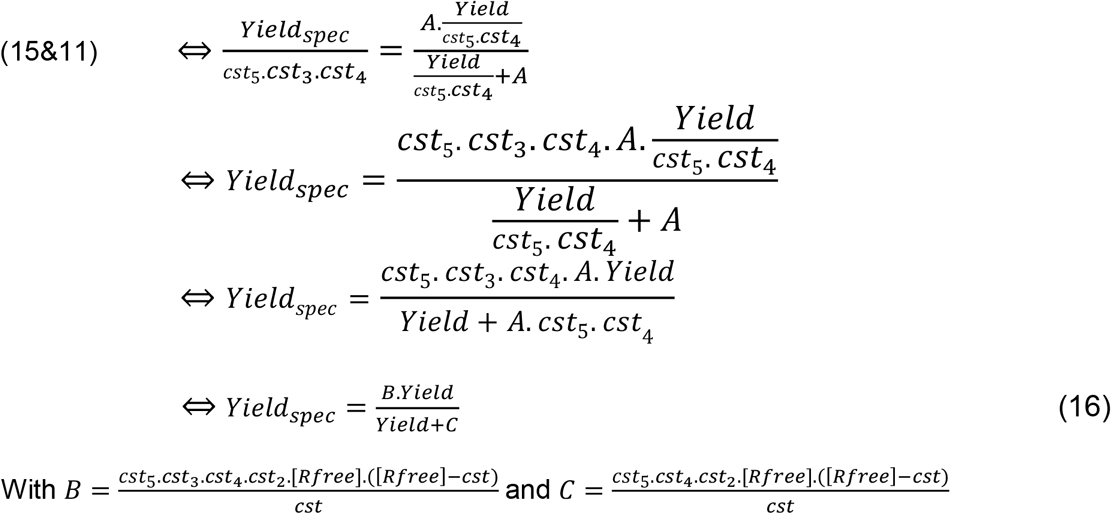

Eventually, we obtained a Michaelis-Menten equation for the relationship between Yield and Yield_spec_ (eq. 16) which explain the data in Supplementary Fig. 6a. Despite the multiple assumptions (that are difficult to verify by experimental measurements) this model gives a simple explanation of our observations.

### SUPPLEMENTARY NOTE 2 Commercial kit and absolute sfGFP measurements (Supplementary Figure 7)

Both plasmids (our plasmid and myTXL plasmid) led to similar yield when the lysate_ORI with the optimized composition (max yield in Fig. 1d) and myTXTL mix are used. This result suggests that pTXTL-P70a(2)-*deGFP* can also be used, instead of our plasmid to optimize cell-free composition. The higher Global yield come from the higher fluorescence obtained with this plasmid. The pTXTL-P70a(2)-*deGFP* seems to be a derivative of the pBEST-OR2-OR1-Pr-UTR1-*eGFP*-Del6-229-T500 ^25^ optimize for expression in cell-free reaction. We don’t have access to the cell-free composition of myTXTL mix but we assumed that it was optimized to obtain a maximum protein production and that the lysate was prepared from a modified strain of *E. coli*. The quality of the result obtained with our lysate-specific optimization compared to the commercial kit is a validation of our method efficiency. The protein concentration obtained from the expression of our plasmid with lysate_ORI is at 0.22 µM sfGFP equivalent. We can notice, with the arbor plasmid, that the the 7 µM sfGFP equivalent is irrelevant as the plasmid produce deGFP.

**Supplementary Figure 1:**
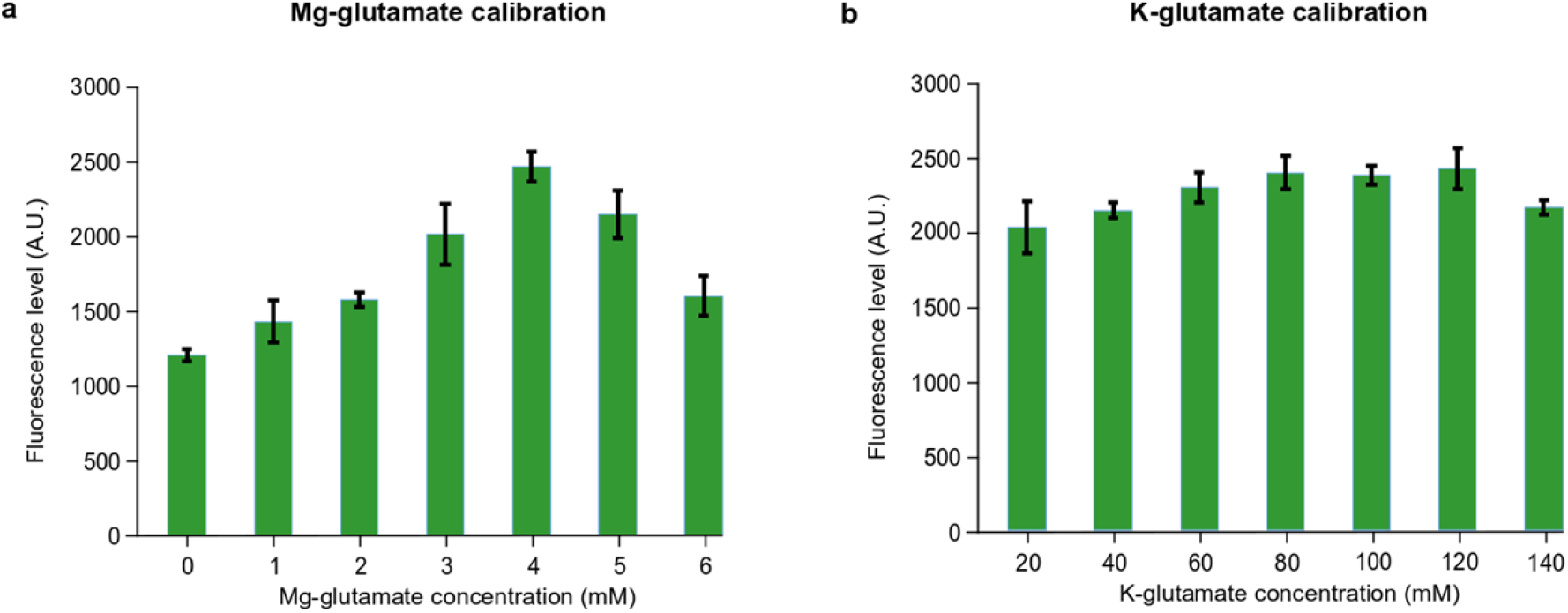
Preliminary calibration of the cell-free composition. The lysate is usually only calibrated for Mg-glutamate, K-glutamate levels. Here we show the end point after overnight cell-free reactions with the lysate_ORI used in Fig. 1. Then, we fixed the maximum concentration for: **a**, Mg-glutamate concentration at 4 mM and **b**, K-glutamate at 80 mM. The error bars stand for the standard deviation of 3 replicates performed on the same day.

**Supplementary Figure 2:**
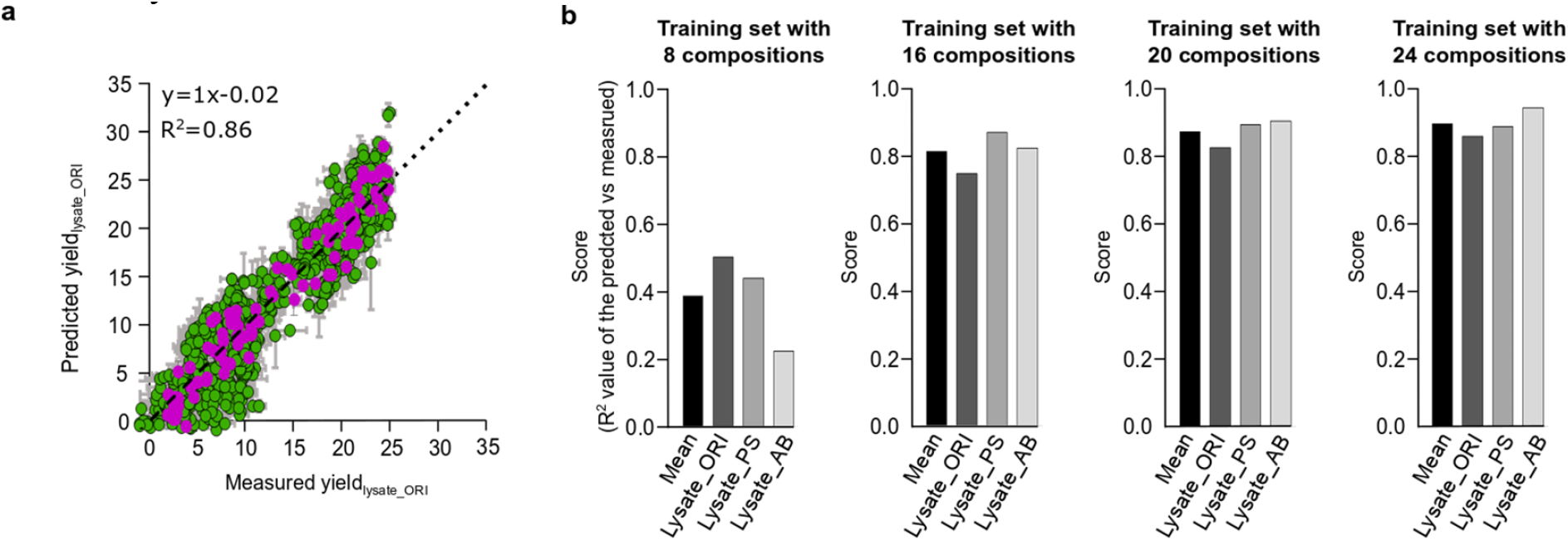
The choice of 102 cell-free compositions for training and testing of our model. **a**, Distribution of the yields obtained with the 102 training cell-free compositions along the 1017 cell-free compositions tested in Fig 1. The 102 cell-free compositions were chosen based on the highest R^2^ obtained by training on 102 points and predicting on the 915 remaining points. The vertical error bars stand for the standard deviation of 3 replicates. The horizontal error bars stand for the standard deviation of 25 predictions. **b**, Comparison of the prediction efficiency of the model when trained with a training set of 8, 16, 20 or 24 cell-free compositions, for prediction on the reminder of the 102 points. The training set is chosen amongst the 102 cell-free compositions fixed in **panel a**. The training set leading to the highest mean R^2^ amongst the 3 lysates has been selected.

**Supplementary Figure 3:**
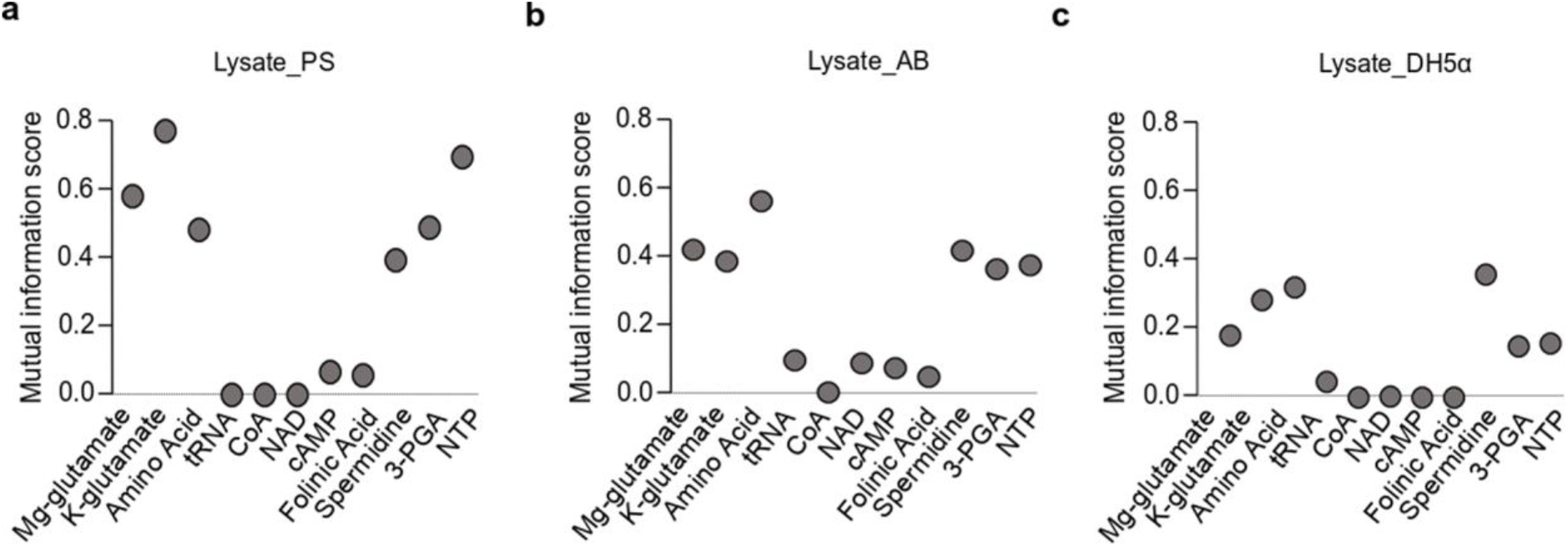
Mutual Information analysis based on the 102 compositions tested with lysate-PS, lysate_AB and lysate_DH5α. Mutual information analysis of the relationship between the yield and each chemical compound, using the yields measured in cell-free reactions using 102 cell-free compositions and **a**, lysate_PS, **b**, lysate_AB, **c**, lysate_DH5α.

**Supplementary Figure 4:**
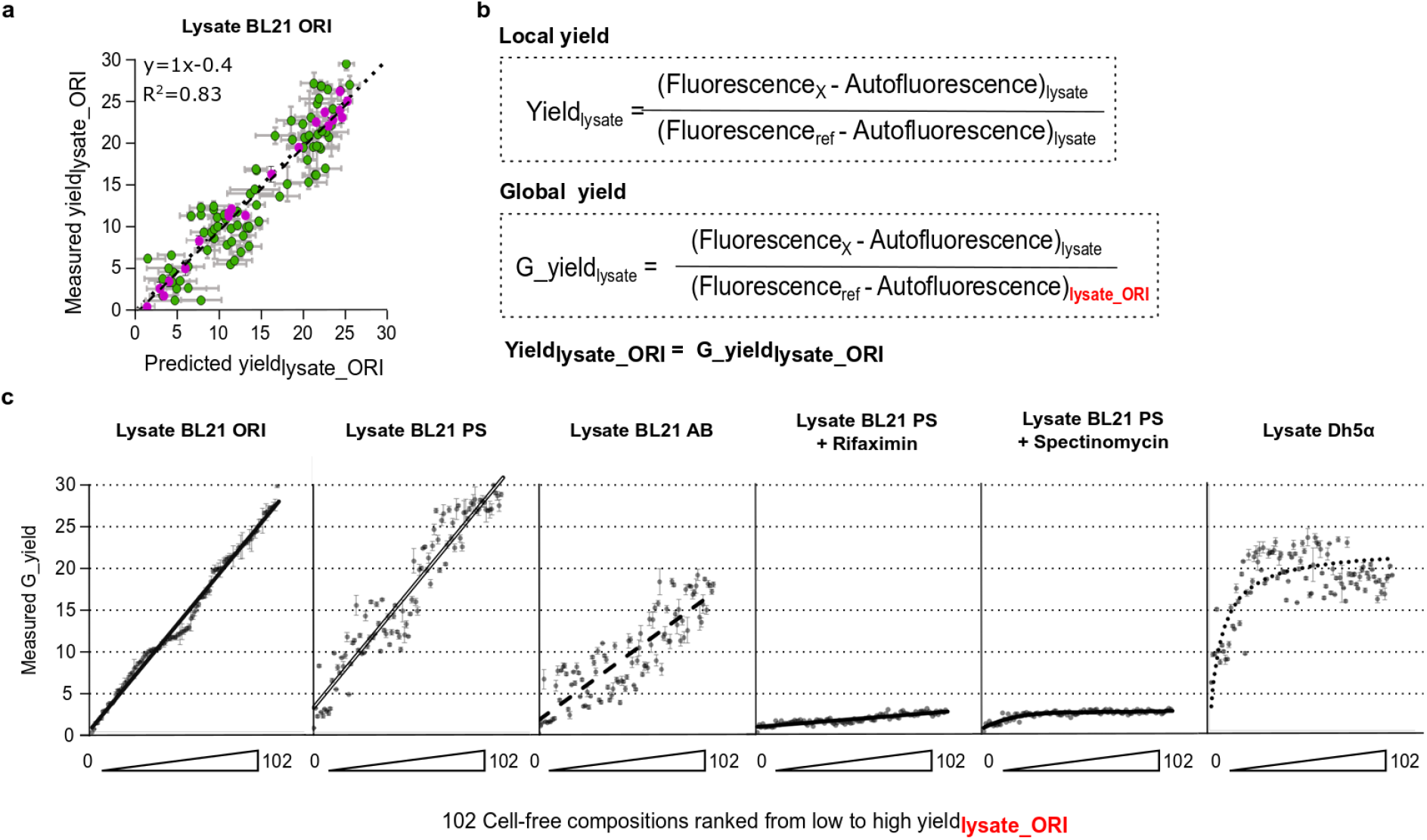
Global comparison between the yields obtained with different lysates. **a**, Comparison of the yields obtained with the lysate original (same as Fig. 1) vs the model predictions for the 102 cell-free compositions used in Fig. 2, Formula of the global yield compared to the local yield. In contrary to the Yields presented in Fig. 2, the Global yield always use the same reference yield from the lysate of Fig. 1 named Lysate_ORI. The Global yield, noted G_yield, allows comparison between yields obtained with our different lysates. **c**, The 102 cell-free compositions were ranked from low to high values based on the yields obtained with the Lysate_ORI. The same ranking of the same 102 cell-free compositions was used for each lysate. Linear fit is used for Lysate_ORI, Lysate_PS, Lysate_AB and Lysate_PS + Riflaximin. Michaelis-Menten like fit is used for Lysate_PS + Spectinomycin and Lysate_DH5α.

**Supplementary Figure 5:**
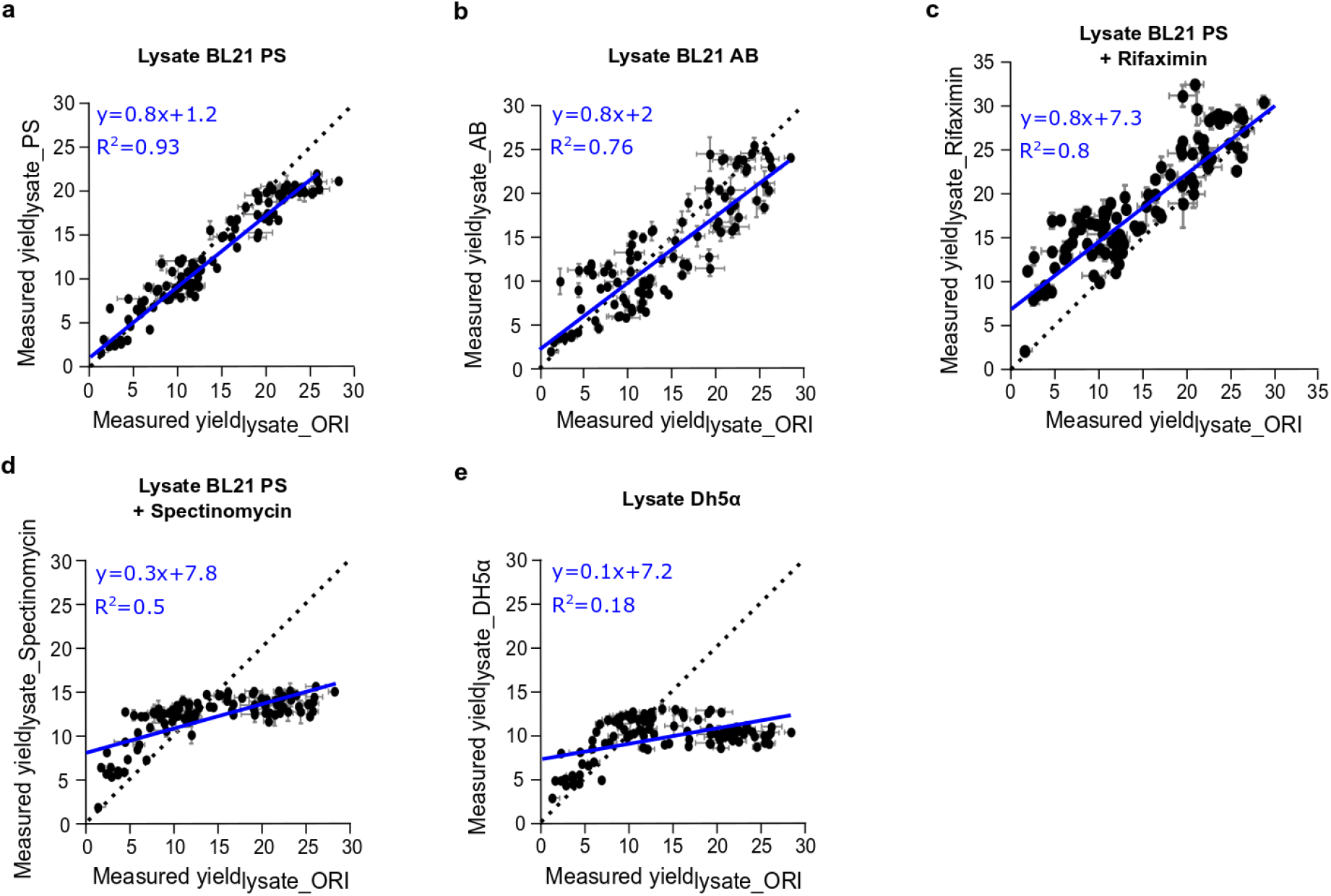
Comparison between the behaviour of the local yields measured with different lysates and the yields measured with the lysate_ORI. Comparison between the yields measured with Lysate_ORI and **a**, Lysate_PS. **b**, Lysate_AB. **c**, Lysate_PS + rifaximin. **d**, Lysate_PS + spectinomycin. **e**, Lysate_DH5α. The blue lines stand for linear fit and the dot lines stand for the perfect correlation (intercept 0 and slope 1). We used the same 102 cell-free compositions for all the measurements. The error bars stand for the standard deviation of 3 replicates

**Supplementary Figure 6:**
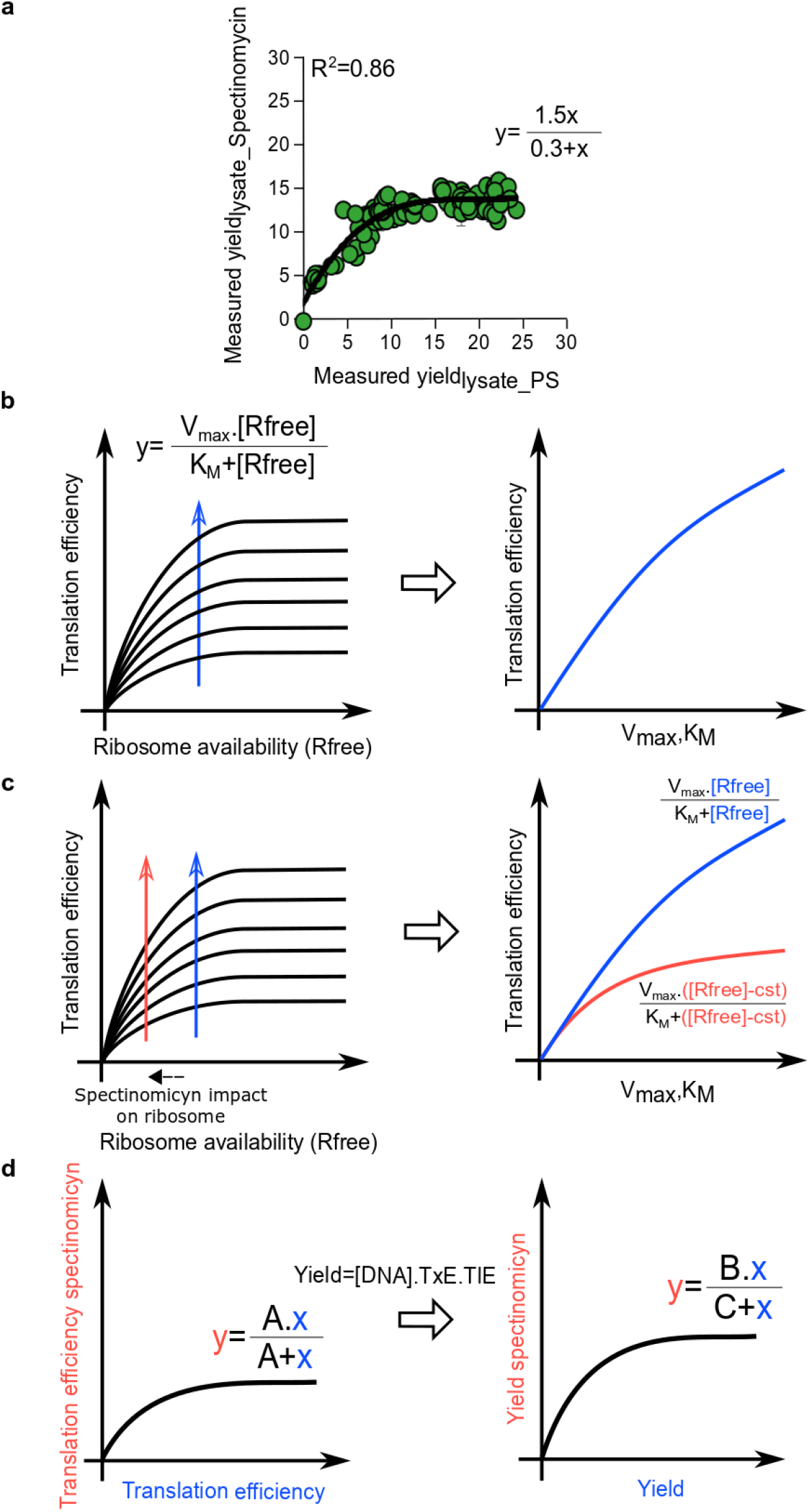
A decrease in ribosome availability is sufficient to explain the saturation of the yields with Lysate_Spectinomycin. **a**, Comparison between the yield obtained with Lysate_PS and the yield obtained with Lysate_PS supplemented with Spectinomycin (same data as Supplementary Fig. 5d). We used a Michaelis-Menten like function to fit the data. **b**, We used the well described Michaelis-Menten^1^ like relationship between translation efficiency and available ribosomes concentration (Rfree). We assumed that a change in cell-composition impact the translation efficiency via a change of V_max_ and K_M_. At a fixed Rfree concentration (blue arrow), an increase of V_max_,K_M_ values lead to an increasing translation efficiency. **c**, As the spectinomycin binds to the 30S subunit of the ribosome to inhibit the translation process, its activity can be represented by a decrease in Rfree concentration (red arrow). The impact of less ribosomes will lead to a decrease in translation efficiency (blue vs red line in the second plot). **d**, Relationship between a translation efficiency with spectinomycin versus a translation efficiency without spectinomycin (see **supplementary note 1**). The yield as the protein production results from the translation but also the transcription process. The relationship between Translation efficiency and yieldsis described in **supplementary note 1**.

**Supplementary Figure 7:**
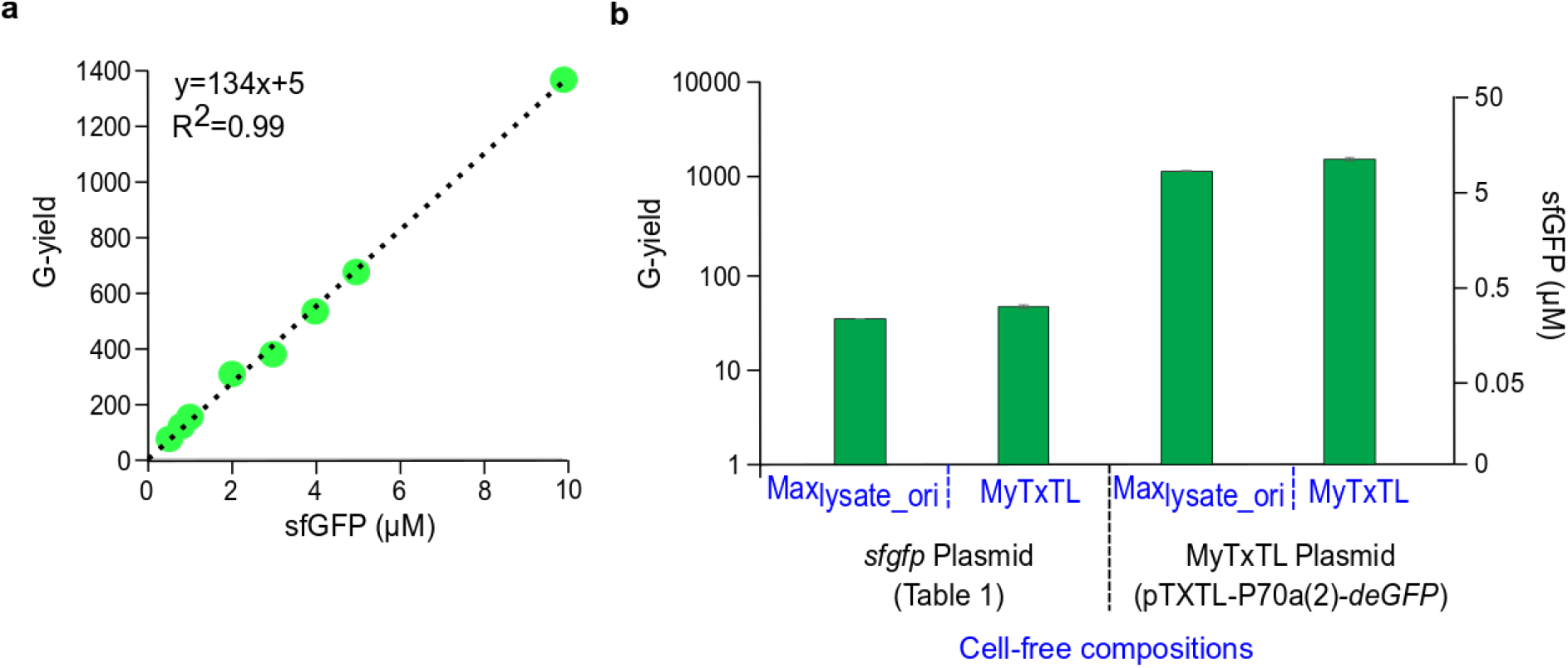
Absolute measurements in cell-free reaction. **a**, Relationship between purified sfGFP and the Global yield. (See **supplementary note 2**). **b**, Comparison between the yield obtained with our best cell-free composition with lysate_ORI and the commercial kit myTXTL from Arbor (pTXTL-P70a(2)-*deGFP*). We used both our plasmid and myTXTL plasmid. Noted that the y-axis with sfGFP concentration is used only for the measurements with our plasmid as myTXTL plasmid produce deGFP and not sfGFP.

**Supplementary Table 1:**
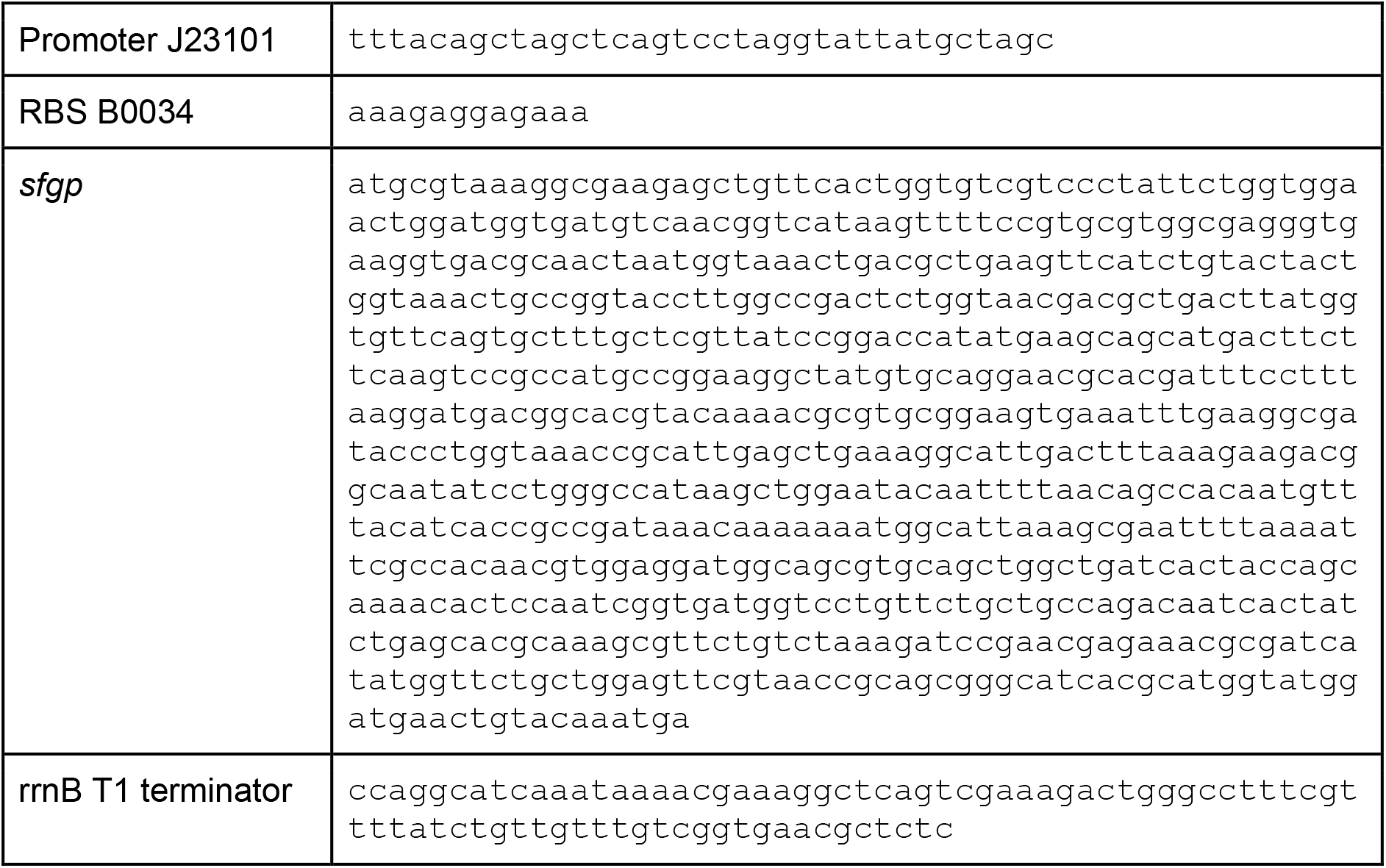
Sequence of the plasmid used in this study. The *sfgp* is under control of the promoter J23101 (http://parts.igem.org/Part:BBa_J23101) and RBS B0034 (http://parts.igem.org/Part:BBa_B0034). The plasmid contains the gene of ampicillin resistance and the origin of replication PBR322.

